# Avoidable false PSMC population size peaks occur across numerous studies

**DOI:** 10.1101/2024.06.17.599025

**Authors:** Leon Hilgers, Shenglin Liu, Axel Jensen, Thomas Brown, Trevor Cousins, Regev Schweiger, Katerina Guschanski, Michael Hiller

## Abstract

Inferring historical population sizes is key to identify drivers of ecological and evolutionary change, and crucial to predict the future of species on our rapidly changing planet. The pairwise sequentially Markovian coalescent (PSMC) method provided a revolutionary framework to reconstruct species’ demographic histories over millions of years based on the genome sequence of a single individual ^1^. Here, we detected and solved a common artifact in PSMC and related methods: recent population peaks followed by population collapses. Combining real and simulated genomes, we show that these peaks do not represent true population dynamics. Instead, ill-set default parameters cause false peaks in our own and published data, which can be avoided by adjusted parameter settings. Furthermore, we show that certain population structure changes can cause similar patterns. Newer methods like Beta-PSMC perform better, but do not always avoid this artifact. Our results suggest testing multiple parameters before interpreting recent population peaks followed by collapses, and call for the development of robust methods.

---

PSMC provides information on past population expansions, contractions, and bottlenecks. To infer changes in effective population size (Ne) over time, PSMC estimates coalescent rates, which are translated into a demographic trajectory assuming neutral evolution under random mating in a single population. PSMC analysis has become a routine analysis for newly-sequenced genomes and has been used to elucidate fundamental questions regarding the evolutionary history of many organisms including humans ^1^, historical drivers of contemporary extinction risk ^2^, and the consequences of climate change ^3,4^. In our investigations of turtle population histories with PSMC, we detected a curious pattern of dramatic peaks followed by even more extreme population collapses (Figure 1 A-F). This pattern occurred across distantly related turtle species around the globe and its existence was consistently supported based on bootstrap replicates. Surprisingly, similar patterns are common in the literature and several studies linked them to climatic or ecological change. For example, an extreme Ne increase followed by a more than 100-fold population collapse in horses around the last glacial maximum was linked to expanding and contracting grasslands ^5^. Similar peaks were further interpreted as indicating the colonization of Southern Africa by Cape Buffalo ^6^, or a transition from initially positive effects of human agriculture to increased hunting pressure on Pink-Footed Goose ^7^. In total, our non-exhaustive search revealed >70 studies with >200 individual PSMC inferences exhibiting extreme peaks followed by population collapses (Supplementary Note). This very pronounced and consistent pattern found in species from across the tree of life raises questions about its biological plausibility and points to a possible presence of analytical artifacts. Such artifacts could cause far-reaching wrong conclusions regarding evolutionary processes, effects of climate change and species conservation.

**Figure 1.**
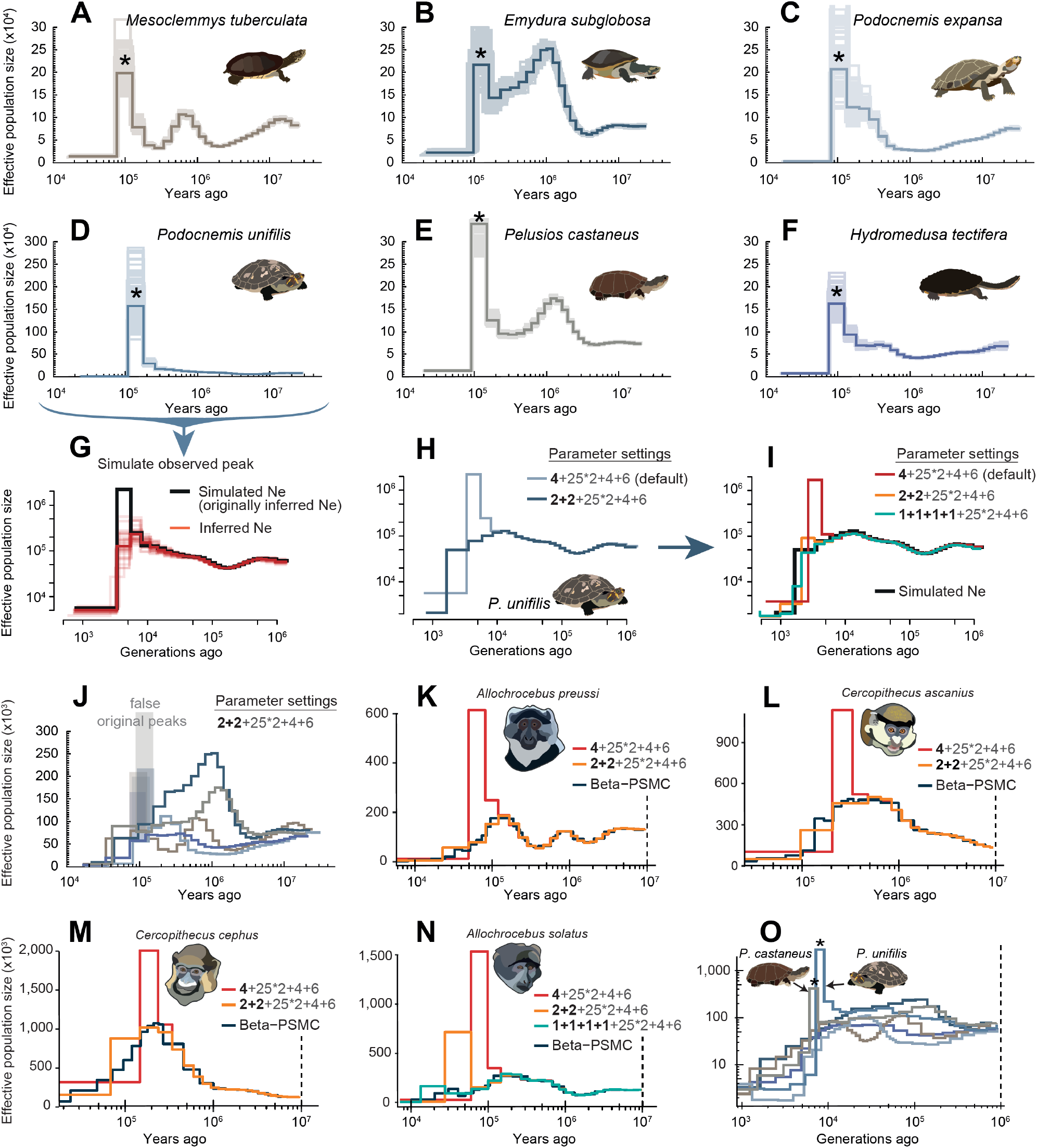
Erroneous peaks with PSMC default settings. **A-F** show effective population size (Ne) inferences for turtles with extreme Ne peaks before population collapses (asterisks) ∼100k years ago. Ne is shown on the y-axis and time in years on the x-axis. Faint lines indicate uncertainty of inferred Ne based on 100 bootstrap replicates. **G** shows mean Ne inference with PSMC (red) based on 30 simulated genomes (pale red) from a population following the Ne trajectory of *P. unifilis* in D (black). **H** shows that splitting the first time window (dark blue) removes the extreme Ne peak inferred in *P. unifilis*. **I** illustrates mean inferred Ne using different parameter settings on 30 simulated genomes with the population history (black) inferred for *P. unifilis* using the split parameter setting (see H). **J** shows that splitting the first time window removes erroneous peaks inferred with PSMC default parameters (faint boxes) in all of our turtle species. **K-N** show effective population size (Ne) inferences for four primate species using several PSMC parameter settings and Beta-PSMC. PSMC default settings recapitulated published peaks (red). Both Beta-PSMC (dark blue) and splitting the first window (yellow and turquoise) removed the erroneous Ne peaks. **O** shows Beta-PSMC results for turtle species using the color code from A-F. Asterisks mark erroneous peaks in two turtles. To aid readability, we excluded very large Ne estimates by Beta-PSMC for the oldest time-windows (dashed lines).

To investigate this, we first asked whether the observed peaks can represent true population size changes. To this end, we simulated diploid genomes (30 replicates) from a population that followed the exact inferred population history of the turtle species with the most pronounced peak in our data: *Podocnemis unifilis* (Figure 1D). Notably, applying PSMC to these genomes, we never detected the extreme Ne peak that we simulated (Figure 1G). This result makes it unlikely that the observed peaks reflect true Ne peaks.

Second, we considered population fragmentation before a population decline as a possible cause. In such a scenario, a single panmictic population would split into subpopulations with limited migration and thus reduced gene flow (Figure S1). Because alleles cannot coalesce as long as they are limited to different subpopulations, the chance of coalescence is reduced during time periods with increased population fragmentation. Since PSMC assumes random mating, reduced coalescence rates caused by population structure will be falsely interpreted as a signature of larger population sizes ^1,4,8^.

To test whether an increase in population structure, not size, could explain the extreme peaks, we used a similar simulation as above, but kept the population size constant before it declines. At the time of the inferred peak in *P. unifilis*, we fragmented the previously panmictic population into three subpopulations with effective migration rates ranging from low (M=0.01) to high (M=10) (Figure S1). Our simulations show that population fragmentation caused a peak that became more prominent with increasing M, approximating the observed original peak at M=5. This shows that certain population structure changes followed by a population collapse can cause patterns resembling the observed peaks. However, it seems unlikely that these processes would always occur in the first and second PSMC time window.

Third, we considered the possibility of a technical artifact related to PSMC parameter settings. Although several studies have used alternate settings, parameters optimized for the human ^1^ are commonly used as default. Most importantly, this default setting fixes the first four atomic time intervals into the first time window. This means PSMC infers a single Ne for a large first time window and cannot model population size changes during this time. For cases with population declines within this window, we reasoned that the model may overcompensate for fixed low Ne by producing exaggerated Ne estimates in the previous time window.

To test this, we applied PSMC to the real *Podocnemis unifilis* data, but split the first time window into two windows (“-p **2+2**+25*2+4+6” instead of the default “-p **4**+25*2+4+6”). In line with our hypothesis, this parameter change completely removes the Ne peak (Figure 1J). To make sure that this behavior is independent of specific genomic features of *P. unifilis*, we simulated 30 genomes that underwent a rapid population decline without a prior radical population size increase, as inferred for *P. unifilis* with split parameter settings (see Figure 1H). Using these genomes, we ran PSMC and found that the artifactual peak was only inferred with the default parameter setting (Figure 1I). This supports that a technical artifact causes erroneous Ne peaks in the second time interval, because a population decline within the fixed first time window cannot be adequately captured by the model. Consistently, splitting the first time window also removed the erroneous peaks in all other turtle species (Figure 1J). Aside from this recent time period, a wide range of settings on both simulated and real data revealed highly congruent results (Supplementary Figure 1D-G).

To confirm that similar peaks in published datasets are also artifacts caused by false parameter settings, we reran PSMC for four primate species exhibiting similar Ne peaks ^9^. As expected, splitting the first time window also removes the peak originally observed for these species (Figure 1 K-N). In one species the population decline reached into the first two atomic time intervals so that the peak only disappeared when the first time window was split into four (-p **1+1+1+1**+25*2+4+6) (Figure 1N).

Since the publication of the original PSMC method, related methods have been developed. However, among the most recent developments, MSMC2 sometimes produces similar extreme peaks ^9^. To test whether other methods also generate erroneous peaks, we ran the recently published beta-PSMC, which should perform better in recent time intervals ^10^. Beta-PSMC outputs for turtle genomes and published primate genomes rarely showed erroneous peaks; however, with a few exceptions (Figure 1 K-O). For the turtles *Podocnemis unifilis* and *Pelusios castaneus*, we also observed erroneous peaks with beta-PSMC (Figure 1O), indicating that this approach does not fully solve the issue.

In conclusion, our results indicate avoidable false population peak inferences across numerous studies, caution against overinterpreting sharp population increases followed by a drastic decline in recent times, and provide a guideline to avoid these artifacts. First, with PSMC, we recommend testing multiple parameter settings that split the first window. Reliable inferences should not be sensitive to different parameter settings. Thus, if a peak is only observed with the default parameter setting, it is likely an artifact. While bootstrapping is used to assess robustness across genomic regions, robustness to different parameters should also be assessed and reported. Second, population structure can be a potential driver of inferred Ne peaks. Third, newer methods such as beta-PSMC avoid many but not all of these artifacts. In addition to the specific tools we investigated, our findings are likely also affect other tools that use PSMC results as input. Our results motivate the development of approaches that solve the problem entirely. For example, it is conceivable to dynamically and automatically split time windows into smaller intervals, as long as enough data per window is available for a robust inference of Ne.

## Acknowledgment

This work was supported by the LOEWE-Centre for Translational Biodiversity Genomics (TBG) funded by the Hessen State Ministry of Higher Education, Research and the Arts (LOEWE/1/10/519/03/03.001(0014)/52). We are grateful to Uwe Fritz, Gene Myers, Peter Praschag, Martin Pippel, Chetan Munegowda and Michail Rovatsos for helpful discussions and their integral roles in the turtle genome project that led to the discovery of the false PSMC peak pattern. We further wish to thank Michael Westbury for helpful discussion and the Bradley Shaffer lab for generating genomes of *M. tuberculata, P. expansa, P. castaneus* and *E. subglobosa* and making the data freely available for analyses. KG was supported by the Swedish Research Council VR (2020-03398), AJ by the Zoologiska Stiftelse grants.

## Author Contributions

Conceptualization, L.H., M.H.; methodology, L.H., S.L., and A.J.; investigation, L.H.; formal analysis, L.H., S.L. and A.J.; writing – original draft, L.H., M.H.; writing – review & editing, L.H., S.L., A.J., T.B., T.C, R.S., K.G., and M.H.; visualization, L.H. and A.J.; funding acquisition, M.H.; resources, M.H.

## Data and Code Availability

All files needed to rerun analyses including the simulated data as well as the code for genome simulations is openly available at figshare.com (DOI: 10.6084/m9.figshare.26975500).

