## Supplement for "Avoidable false PSMC population size peaks occur across numerous studies"

### Figure S1

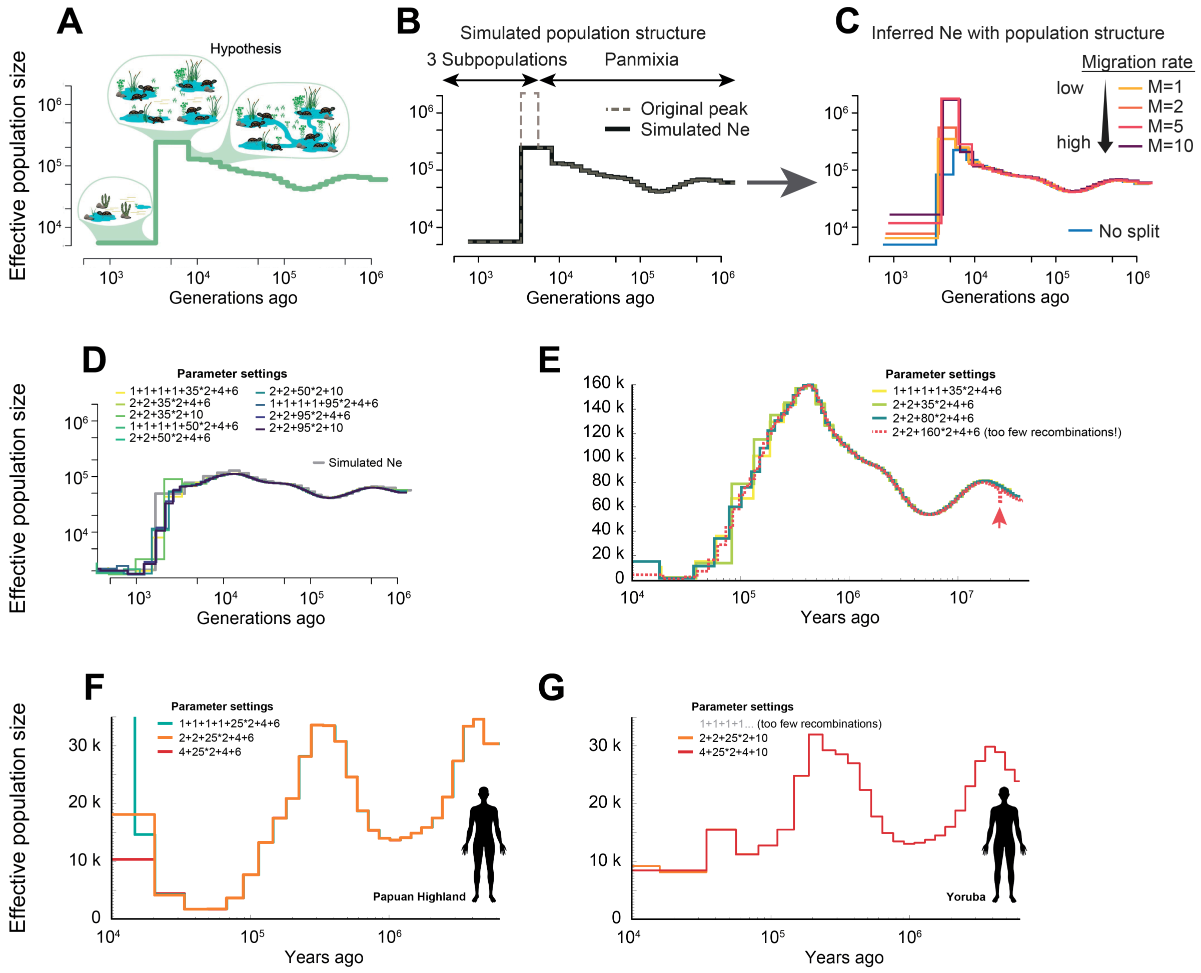


**Figure S1: Inferring Ne with different parameter settings and changes in population structure.** **A** illustrates how climate change, e.g., increasing aridity, may cause increased population structure resulting in the formation of subpopulations, before a population collapse. This scenario may cause reduced gene flow between subpopulations in the plateau phase resulting in reduced coalescence rates and false inference of increased Ne using PSMC. **B** illustrates a simulation of the Ne trajectory. Instead of a true Ne peak, a panmictic population splits into three subpopulations. Mean inferred Ne based on 30 individual genomes from this simulation with different migration rates are shown in **C**. While our data show that such a scenario can result in erroneous peaks, the most likely and most common cause for these peaks are likely false parameter settings (see main text). Mean inferred Ne using a wide range of different parameter settings based on 30 genomes that evolved following the trajectory shown in grey is displayed in **D**. **E** shows inferred Ne for *Podocnemis unifilis* using a wide range of different parameter settings. A red arrow shows small signs of overfitting when too many time windows are introduced. Generally, all tested parameter settings are in high agreement with each other regarding the deeper inferred population history using either real and simulated data. Hence, the only main exception is the artifact for the recent history caused by the default parameters reported in the main text of this manuscript. The deeper demographic history can be robustly inferred with either the default or other, more fine-grained, parameter settings as long as enough recombinations are inferred for each time window. Human population sizes for Papuan Highlands and Yoruba are shown in **F** and **G**, respectively. Here, inferences using different parameters are highly congruent and only differ for very recent time periods less than 20k years ago.

### Supplemental Experimental Procedures

#### Read mapping and variant calling

For turtles, 10X reads were mapped to published genomes of *M. tuberculata* (GCA_007922155.1), *P. expansa* (GCA_007922195.1), *E. subglobosa* (GCA_007922225.1) and *P. castaneus* (GCA_007922175.1), as well as newly generated high-quality chromosome-level reference genomes of *H. tectifera* and *P. unifilis* (BUSCO completeness >99.5%, Contig N50 > 30MB) using Longranger (v2.2.2). Specifically, the longranger align command was used to group reads based on the 10X barcodes, trim these barcodes from the input sequences, perform error-correction, sort and align the reads to the genomes. Resulting mappings were quality adjusted (mpileup -C50), filtered to remove sites with less than one third or more than twice the average genome wide coverage with bcftools (v1.10.2) [S1]. Mean coverages raged from 48X to 85X (*E. subglobosa*: 48X, *M. tuberculata*: 54X, *P. castaneus*: 52X, *P. unifilis*: 85X, *P. expansa*: 48X, *H. tectifera*: 76X). Finally, variant calls (including invariant sites) were converted to fasta sequences and the PSMC-specific input format using fq2psmcfa (-q20) (https://github.com/lh3/psmc).

The primate genomes (Illumina paired end 150 bp, ~30X coverage) were processed following the GATK best practices for germline variant discovery [S2]. Briefly, we aligned the reads to the macaque (Macaca mulatta) reference genome (Mmul_10, GCF_003339765.1), filtered out PCR-duplicates with Picard/2.23.4 and called variants with GATK/4.2 HaplotypeCaller and GenotypeGVCFs, set to output also invariant sites. Variants were filtered with GATK VariantFiltration using the recommended exclusion criteria (QD < 2.0, QUAL < 30.0, SOR > 3.0, FS > 60.0, MQ < 40.0, MQRankSum < -12.5, ReadPosRankSum < -8.0). We also removed sites where read coverage was less than half or more than twice the genome wide average, and masked repetitive regions following the SNPable regions pipeline (https://lh3lh3.users.sourceforge.net/snpable.shtml). Filtered autosomal variant calls (including invariant sites) were converted to fasta sequences and the PSMC-specific input format psmcfa using bcftools [S1] and fq2psmcfa (https://github.com/lh3/psmc), respectively.

High quality human data (Samples: HGDP00551, HGDP00931) (Figure S1 F, G) was downloaded from the 1000 human genomes project (https://ftp.1000genomes.ebi.ac.uk/vol1/ftp/data_collections/HGDP/data/) and .cram files were converted to .bam files using samtools v1.14 [S1]. Subsequently, data was processed identically to the turtle dataset.

#### Historical population size inference using PSMC and Beta-PSMC

We ran PSMC with the settings "-N25 -t15 -r5", varying the parameter patterns (-p) between "4+25*2+4+6", "2+2+25*2+4+6" and, if needed, "1+1+1+1+25*2+4+6". To further test the effect of more parameters in deeper time intervals, we ran a broad range of parameters including up to 164 time intervals on simulated and real data (see Figure S1 D, E). We used splitfa from the psmc-utils to split the autosomal psmcfa into shorter segments from which bootstrap replicates were subsequently sampled. Twenty and one hundred bootstrap replicates were run for primates and turtles, respectively. To avoid overfitting, we manually confirmed that at least 10 recombination events were inferred for each time window after 20 iterations. The only exception to this is when we specifically tested the effect of potential overfitting by setting parameters to "-p 2+2+16*2+4+6" on *P. unifilis* (Figure S1 E). For turtles, PSMC inferences were scaled using a mutation rate of $7.9 \times 10^{-9}$ [S3, S4] and generation times of 15 years for *Emydura subglobosa* and *Pelusios castaneus*, 25 years for *Podocnemis unifilis*, *Hydromedusa tectifera* and *Mesoclemmys tuberculata*, and 50 years for *Podocnemis expansa*. The primate PSMC inferences were scaled using a mutation rate of $4.91 \times 10^{-9}$ and a generation time of 10 years for *Allochrocebus solatus* and *A. preussi*, and a mutation rate of $4.82 \times 10^{-9}$ and generation times of 8 and 12 years for *Cercopithecus cephus* and *C. ascanius*, respectively [S5]. Human inferences were scaled using a mutation rate of $1.25 \times 10^{-8}$ and a generation time of 25 years. For Yoruba, the last two time intervals were merged to achieve enough recombination events per time window (Figure S 1G).

Beta-PSMC extends the original PSMC model by allowing the population to vary within each discretized time interval [S6]. As this improves resolution, especially in recent times, we explored whether Beta-PSMC was less prone to produce erroneous Ne spikes. Following the authors recommendations, we ran Beta-PSMC only with 20 parameter vectors spanning one discretized time interval (-p 20*1).

**Simulated genomes**
We used "psmc2history.pl" and "history2ms.pl" to convert the PSMC outputs to an ms command. For each analysis we then simulated genomes (30 replicates) with 10 pairs of 30 Mb chromosomes using coalescent simulation with recombination [S7].

**Published peak examples**
We carried out a non-exhaustive search for published PSMC (and related methods like MSMC2) results with peaks in the second time interval, followed by a population collapse to very low Ne in the first time interval. Examples of studies reporting such results are listed [S5, S8-S78].
